# Diffusion of metabolites across gap junctions mediates metabolic coordination of β-islet cells

**DOI:** 10.1101/2020.12.23.424180

**Authors:** Vishnu P. Rao, Megan A. Rizzo

## Abstract

Loss of oscillatory insulin secretion is an early marker of type II diabetes. In an individual beta-cell, insulin secretion is triggered by glucose metabolism, which leads to membrane depolarization and calcium influx. Islet β-cells display coordinated secretion; however, it is unclear how the heterogeneous population of insulin-secreting beta cells coordinate their response to glucose. The mechanisms underlying both electrical and calcium synchronicity are well explored. Even so, the mechanism governing metabolic coordination is unclear given key glycolytic enzymes’ heterogeneous expression. To understand how islet cells coordinate their metabolic activity, a microfluidic applicator delivered glucose stimulation to spatially defined areas of isolated mouse islets. We measured metabolic responses using NADH/NADPH (or NAD(P)H) autofluorescence and calcium changes using Fluo-4 fluorescence. Glucose stimulated a rise in NAD(P)H even in islet areas unexposed to the treatment, suggesting metabolic coordination. A gap junction inhibitor blocked the coordinated NAD(P)H rise in the low-glucose areas. Additionally, metabolic communication did not occur in immortalized β-cell clusters known to lack gap junctions, demonstrating their importance for metabolic coupling. Metabolic communication also preceded glucose-stimulated rises in intracellular calcium. Further, pharmacological blockade of calcium influx did not disrupt NAD(P)H rises in untreated regions. These data suggest that metabolic coordination between islet beta-cells relies on gap-junctional activity and precedes synchronous electrical and calcium activity.

## Introduction

The inability of glucose-stimulated insulin secretion to compensate for peripheral insulin resistance contributes to type 2 diabetes progression^1–3^. Since cellular communication within an islet potentiates secretory output tenfold^4–6^, intra-islet signaling mechanisms provide vital information about secretory failure and treatment strategies in diabetes. Therapeutic approaches to restore or enhance insulin secretion are viable both experimentally^7–9^ and clinically^10,11^. As such, it is beneficial to understand how intra-islet signaling pathways promote insulin secretion.

Within individual beta cells, the glucose-stimulated insulin secretion pathway is well defined. Fluctuations in blood glucose proportionally alter cellular metabolism via changes in glucokinase activity^12,13^. Sufficient change in the ATP/ADP ratio, in turn, triggers an action potential, calcium influx, and fusion of insulin granules with the plasma membrane^14^. However, islet insulin secretion, specifically the secretory boost that arises from the organization of beta cells in islets, is incompletely understood. Beta cells, even within an individual islet, are heterogeneous in important ways. They have different glucose stimulation thresholds for metabolic activity and insulin secretion^15^ as well as distinct gene and protein expression^16–20^. For example, glucokinase expression is increased in high insulin secreting beta cells^21^ while beta cells with increased expression of genes encoding ATP-sensitive potassium channel (KATP) subunits display lower insulin secretion^22^. Despite this heterogeneity, islet beta cells respond in unison. NAD(P)H levels rise similarly after glucose treatment^23,24^, suggesting that cell-cell communication occurs before membrane depolarization. Action potential dynamics are also shared across all islet beta cells leading to synchronous calcium oscillations^25^.

Yet, the mechanisms underlying synchronous islet beta cell responses are unresolved^26^. Gap junctions connecting cells are crucial for coordinating intra-islet beta cell activity^27,28^. Chemical inhibition of gap junctional coupling or knock out of proteins forming gap junctions results in loss of islet calcium oscillations and pulsatile insulin secretion^27–29^. Although a wide range of studies^30–35^ have established the prominence of gap junctions, it is unclear which molecules coordinate the cell-cell communication in the islet. Metabolites^36,37^, nucleotides^38^, and ions^39^ have each been proposed. Electrical coupling is the most straight forward to test and best described^40,41^. Even so, models based on electrical coupling due to ionic diffusion best account for cell-cell coordination during an action potential. Beyond establishing an islet-wide threshold potential, ionic flow through gap junctions would have little impact on pre-threshold cell-cell communication.

Metabolites are logical mediators of pre-action potential communication through gap junctions, although direct evidence has proved elusive. In theory, islet gap junctions are large enough pores to pass several interesting cell-signaling candidates. Lucifer yellow, for example, freely passes through islet gap junctions^42,43^. At 442 Da, it is larger than notable cell signaling intermediates including inositol trisphosphate (420 Da, charge = −2 to −4) and cAMP (329 Da, charge = −1). Glucose 6-phosphate, fructose 1,6-bisphosphate, and ADP can also diffuse across monolayer beta cells from neonatal islet monolayer cultures^36^. Given the notable differences between monolayer culture and naturally formed adult islets^44–46^, whether metabolic diffusion occurs between beta cells in intact islets has not been sufficiently explored. In this study, we clarify the presence of metabolic coupling using selective stimulation of islets. Delivery of glucose to spatially defined areas of adult rodent islets reveals metabolites diffuse across gap junctions independent of rises in cytoplasmic calcium. These results implicate metabolic coupling in coordination of islet secretory responses and maintenance of appropriate insulin secretion.

## Results

Previous studies have shown that metabolic responses to glucose are similar across the islet following glucose stimulation^23,47,48^. Typically, the entire islets’ bathing solution receives additional glucose using a perfusion system or manual pipetting^49–52^. To determine if metabolic responses spread across beta cells, we used a microfluidic approach to stimulate a portion of the adult mouse islet. We then measured metabolic responses in stimulated and unstimulated regions. The FluiCell Biopen Prime system administered glucose using positive and negative pressures to restrict chemical exposure to a localized volume^53,54^. NAD(P)H autofluorescence, a marker of metabolic activity^55–57^, was used to quantify metabolic responses. Before glucose simulation, we sought to identify islet portions to which the solution was delivered (stimulated regions) and restricted from (unstimulated regions). Areas in the treated zone were marked using rhodamine 101. The area exposed to the dye was visually distinct (Figure 1A). Using an image thresholding algorithm, stimulated and unstimulated regions were determined and match what is visually observed (Figure 1B). Further, the islet and stimulation edges were computationally identified (Figure 1C). These metrics allowed us to quantify islet responses based on distance from the islet edge or the stimulation edge in a pixel-by-pixel manner.

**Figure 1.**
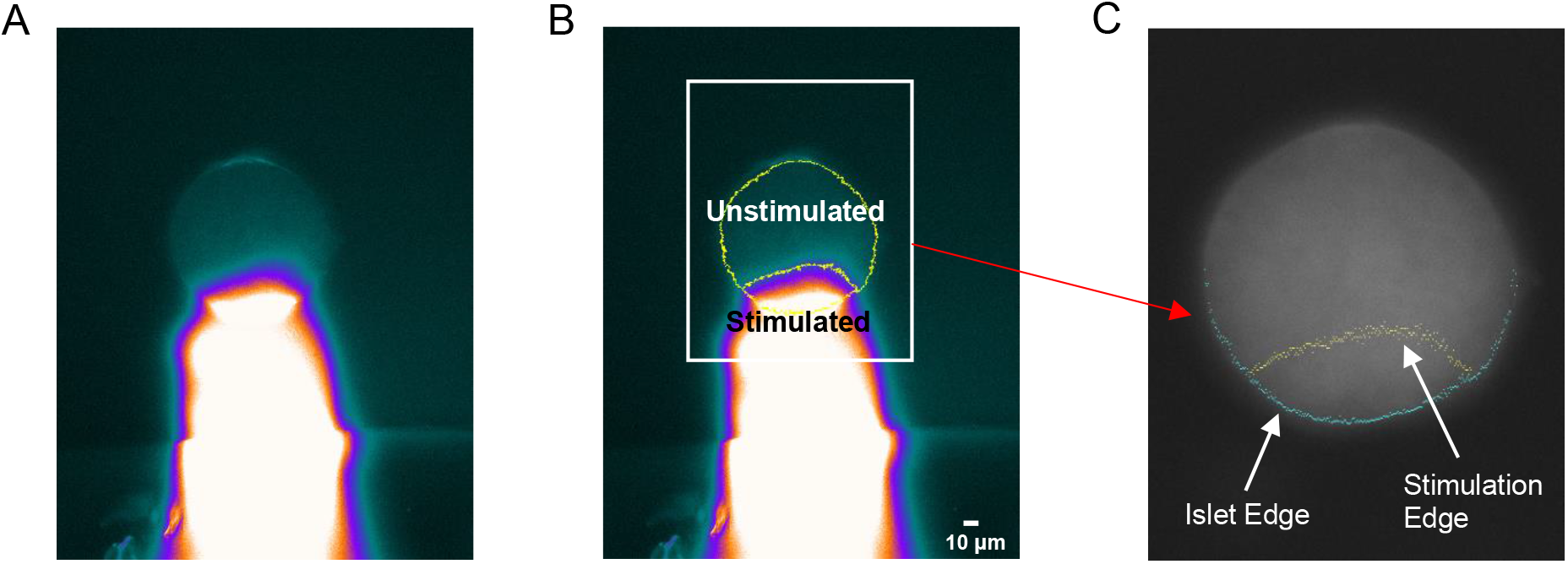
Identification of stimulated and unstimulated islet regions. (A) Regions of the islet were exposed to 500 nM rhodamine 101 dye using a microfluidic applicator. (B) A thresholding algorithm identifies exposed (stimulated) and unexposed (unstimulated) islet regions, outlined in yellow. A magnified view of the islet (white rectangle) is shown in (C). The border of the islet facing the microfluidic applicator (islet edge, blue) as well as the boundary to which solution is dispensed (stimulation edge, yellow), as indicated by the white arrows, are determined computationally.

After delineating stimulated and unstimulated regions, we tested whether the solution delivered by the microfluidic applicator was indeed confined to a localized volume. The INS-1E glucose responsive rat insulinoma cell line lacks gap junctions^58,59^. Thus, metabolic responses should be appropriately confined to regions stimulated by glucose. Stimulated and unstimulated regions were determined, as in Figure 1, with rhodamine dye. Treatment with 20 mM glucose from a resting 2 mM glucose revealed that responses are restricted to stimulated regions in INS-1E clusters (Figure 2). Average NAD(P)H responses over time show rises in stimulated regions that were not detected outside of the rhodamine-marked area (Figure 2A). Control treatment with 2 mM glucose, to assess the effect of fluid force, shows only a slight increase in NAD(P)H that is distinct from glucose-dependent rises (Figure 2B). A visual representation of NAD(P)H responses is depicted for an INS-1E cluster exposed 20 mM glucose (Figure 2C). The pseudocolor map of responses illustrates responses are confined to the exposed region, outlined in yellow. Another indication that the delivery of solution is constrained is based on the max NAD(P)H response across distance. The INS-1E cell line is a relatively homogeneous, clonally derived beta cell line^60^. Therefore, NAD(P)H responses should be equal throughout portions exposed to glucose. In stimulated regions, NAD(P)H responses are even with a non-significant slope (p=0.6261) across distance (Figure 2D). The same holds true for unstimulated regions (p=0.1296) (Figure 2E). Importantly, max NAD(P)H responses are significantly greater (p<0.0001) in stimulated regions of INS-1E clusters (Figure 2F). Thus, only the area marked by rhodamine is stimulated by the microfluidic applicator.

**Figure 2.**
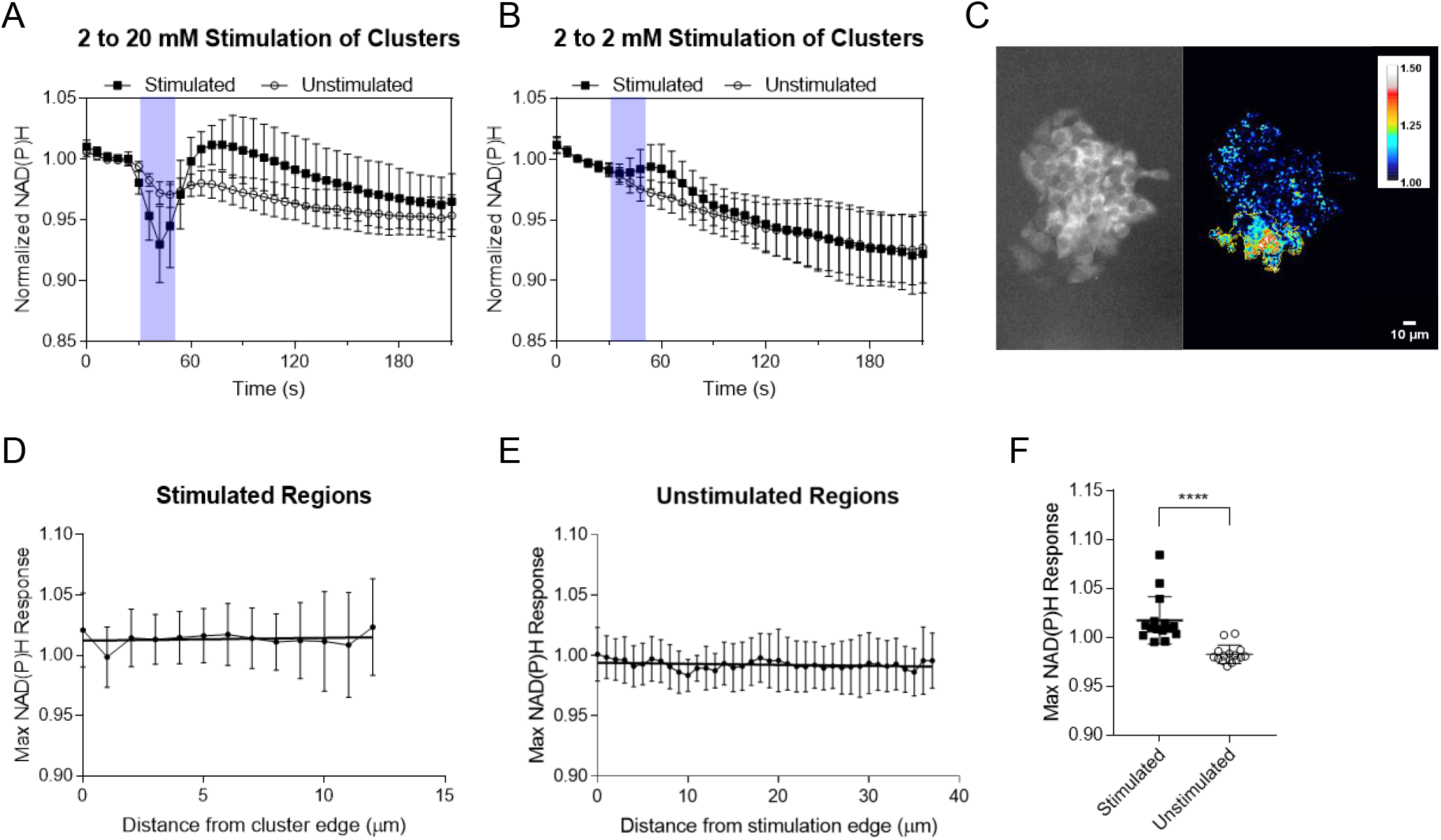
Metabolic responses are confined to stimulated regions in INS-1E clusters. (A) INS-1E clusters were exposed to 20 mM glucose for 20 seconds, purple bar, using a microfluidic applicator. Mean NAD(P)H responses (symbols) ± S.D. (error bars) in stimulated and unstimulated regions are displayed (n=15 clusters). Stimulated regions show a response to glucose while unstimulated regions do not. (B) 2 mM glucose was applied, purple bar, to clusters as a control to observe the effect of pressure. A small rise in NAD(P)H is seen in stimulated regions. Mean NAD(P)H responses ± S.D. are depicted (n=15 clusters). (C) A pseudocolor map of NAD(P)H responses following administration of 20 mM glucose is shown for an INS-1E cluster. The stimulated region is outlined in yellow. Metabolic responses are restricted to the stimulated region. (D, E) NAD(P)H responses were averaged at integer distances (n=15 clusters). The maximum, across time, of these average responses is plotted versus distance (error bars indicate S.D.). Metabolic responses are consistent and do not correlate with distance in stimulated regions (p=0.6261, r^2^=0.02232) or unstimulated regions (p=0.1296, r^2^=0.06266). (F) Max NAD(P)H responses are greater in stimulated regions compared to unstimulated regions (n=15 clusters, p<0.0001). Symbols denote individual islets. Linear regression was used to evaluate response versus distance relationships and a matched pairs t-test was used to compare max NAD(P)H responses.

Next, we explored if islets are metabolically coordinated, i.e., whether metabolic responses spread between cells. We started by treating portions of islets with 11 mM glucose from a resting 5.5 mM glucose, which are physiologic postprandial and fasting blood glucose concentrations. However, given the short duration of treatment, it was hard to detect metabolic responses. NAD(P)H responses marginally rise above baseline in stimulated regions and cannot be observed in unstimulated regions (Figure 3A). To accentuate metabolic responses, we moved from a two-fold increase (5.5 to 11 mM) to a ten-fold increase (2 to 20 mM). With the ten-fold treatment, above baseline NAD(P)H responses are observed in both stimulated and unstimulated regions (Figure 3B). Metabolic responses appear to occur later and decrease in magnitude in areas further from the stimulation edge (Figure 3C-D). Similar to INS-1E clusters, treatment with 2 mM glucose shows slight rises in NAD(P)H (Figure 3E). To examine how max NAD(P)H responses change with distance, we used a pixel-by-pixel analysis to quantify the response. In stimulated regions of the islet, maximum responses slightly increase with distance from the islet edge (Figure 4A). Interestingly, maximum responses in unstimulated areas correlate well with distance, decreasing linearly as distance from the stimulation edge increases (r^2^=0.8639, p<0.0001) (Figure 4B). These relationships also hold true for responses at a given timepoint (Supplemental Figure 1). We then compared the time to the maximum response at the pixel level. The peak response in unstimulated areas occurred significantly later (p=0.0027) than in regions directly exposed to glucose. This result suggests that metabolic responses spread throughout the entire islet and are not constrained to cells directly exposed to glucose (Figure 4C). Given that responses decay linearly with distance in unstimulated regions, these data point to a coordinating molecule’s unhindered diffusion.

**Figure 3.**
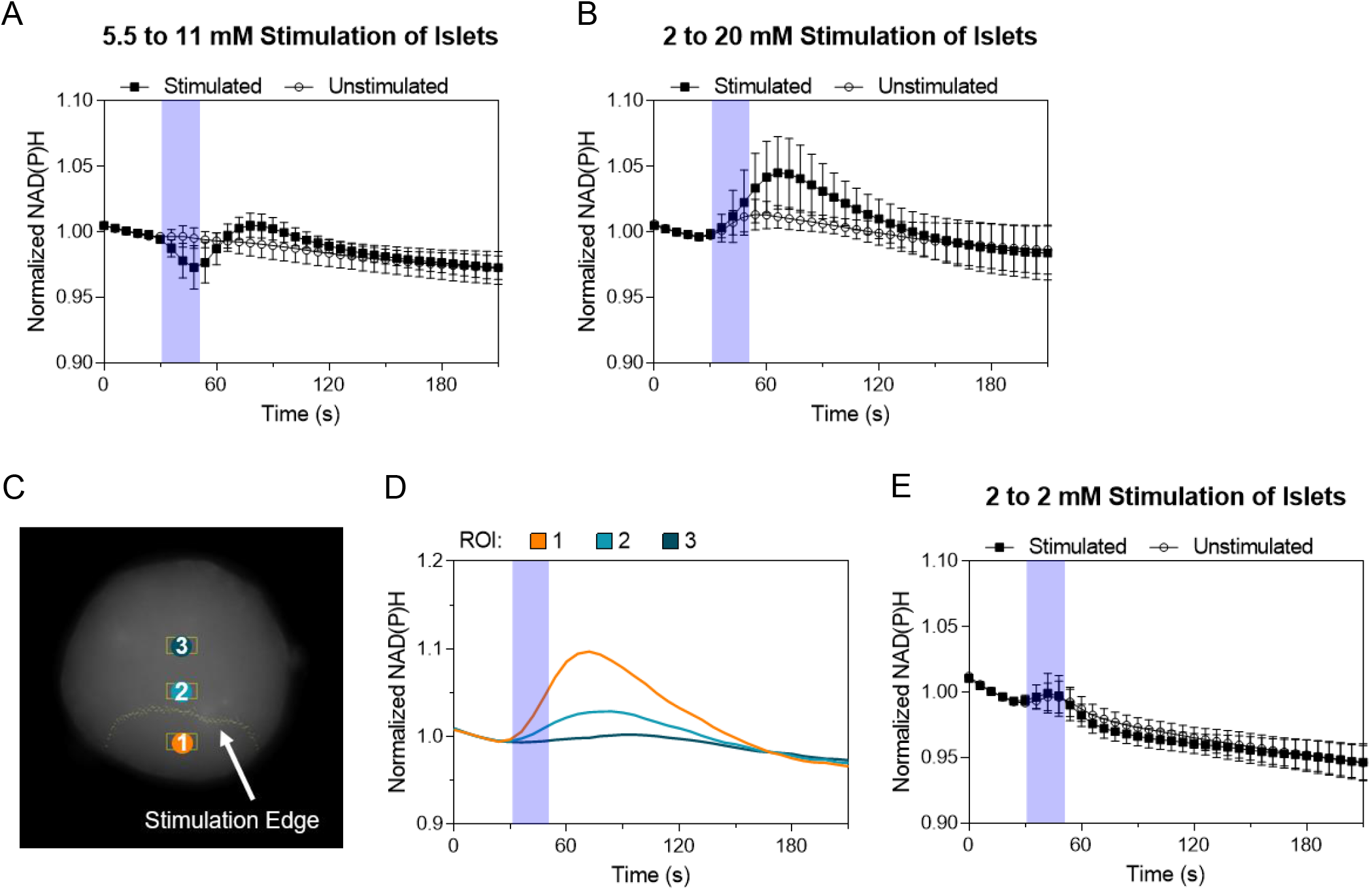
Metabolic coordination exists in islets. (A) Islets were exposed to 11 mM glucose for 20 seconds, purple bar, from a baseline 5.5 mM glucose using a microfluidic applicator. Small rises in NAD(P)H are observed in stimulated regions, but no such rises are seen in unstimulated regions of the islet. Mean NAD(P)H responses (symbols) ± S.D. (error bars) are presented (n=16 islets). (B) Application of 20 mM glucose for 20 seconds, purple bar, from a baseline 2 mM glucose shows NAD(P)H responses in stimulated and unstimulated regions (n=16 islets). (C) An example islet is labelled with three regions of interest (yellow boxes and colored circles) and the stimulation edge (yellow, indicated by white arrow). (D) NAD(P)H responses, from each region of interest (ROI), across time is displayed. Responses decrease in magnitude and occur later in ROIs beyond the stimulation edge. (E) Stimulation with 2 mM glucose, purple bar, from a baseline 2 mM glucose shows a slight rise in NAD(P)H that is distinct from glucose dependent responses (n=16 islets).

**Figure 4.**
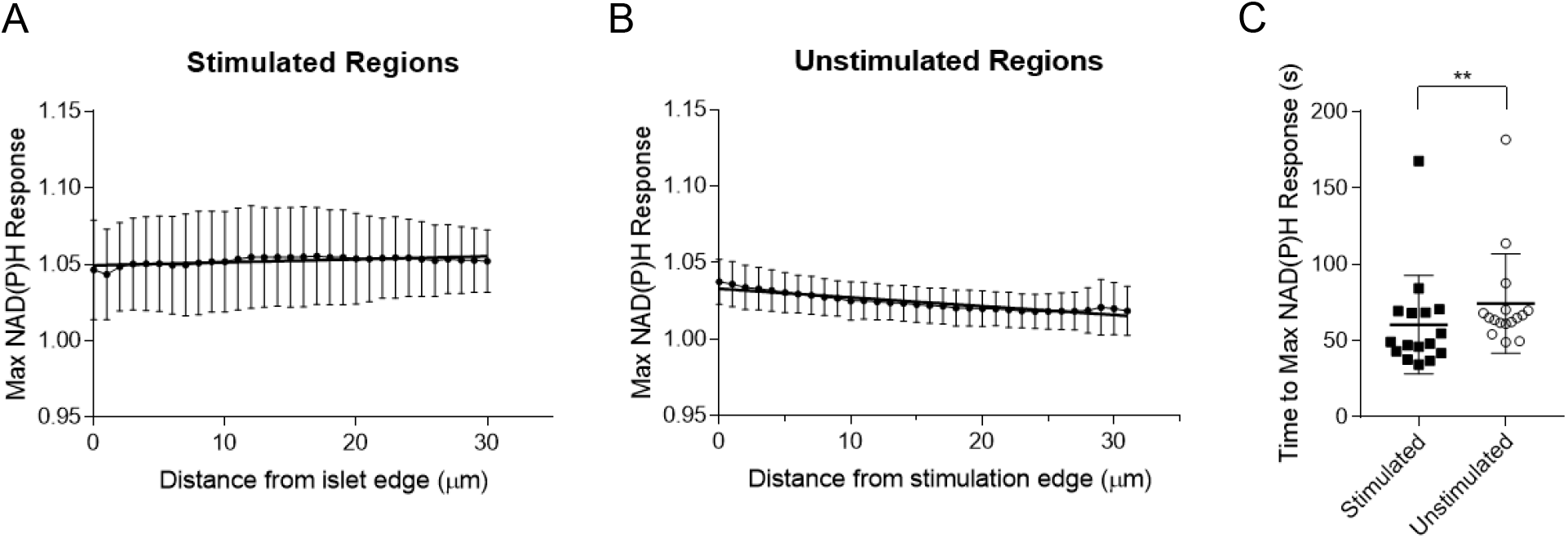
Metabolic responses are consistent with diffusion of a coordinating molecule. Portions of islets were exposed to 20 mM glucose from a baseline 2 mM glucose using a microfluidic applicator. (A, B) NAD(P)H responses were averaged at integer distances (n=16 islets). The maximum, across time, of these average responses is plotted versus distance (error bars indicate S.D.). Responses in stimulated regions (A) slightly rise with distance from islet edge (p<0.0001, r^2^=0.431, slope=0.0001986 ± 4.238e-005) while responses in unstimulated regions (B) decrease and correlate well with distance from islet edge (p<0.0001, r^2^=0.8639, slope= −0.000569 ± 4.124e-005). (C) The time to max NAD(P)H response following glucose treatment is shown for stimulated and unstimulated regions from corresponding islets (n=16 islets). Responses in stimulated regions peak significantly earlier than responses in unstimulated regions (p=0.0027). Response versus distance relationships were assessed by linear regression. The difference in the time to max NAD(P)H responses was evaluated by Wilcoxon matched-pairs signed rank test.

Although the spread of metabolic responses indicates diffusion, we wanted to test whether gap junctions mediated this. We treated islets with an inhibitor of gap junctional communication, carbenoxolone^61–64^, before glucose application (Figure 5). In the presence of carbenoxolone, responses are abolished only in unstimulated regions (Figure 5A). We also looked at the difference in response between the stimulated and unstimulated regions. Compared to untreated islets, islets treated with carbenoxolone do not show a greater difference in response (p=0.1985) (Figure 5B-C). However, responses still follow a similar time course as the time to max response is unchanged (p=0.3608) (Figure 5D). To measure off-target effects, we treated INS-1E clusters, which lack gap junctions, with carbenoxolone before administering glucose. NAD(P)H responses in both regions are unchanged (stimulated regions: p=0.8215, unstimulated regions: p=0.2706) with carbenoxolone treatment (Figure 5E-F). Like the untreated clusters in Figure 2, responses are significantly greater and only observed in stimulated regions compared to unexposed areas (data not shown, p=0.0353). We can conclude that gap junctional diffusion of some molecule(s) coordinates the metabolic responses from these data.

**Figure 5.**
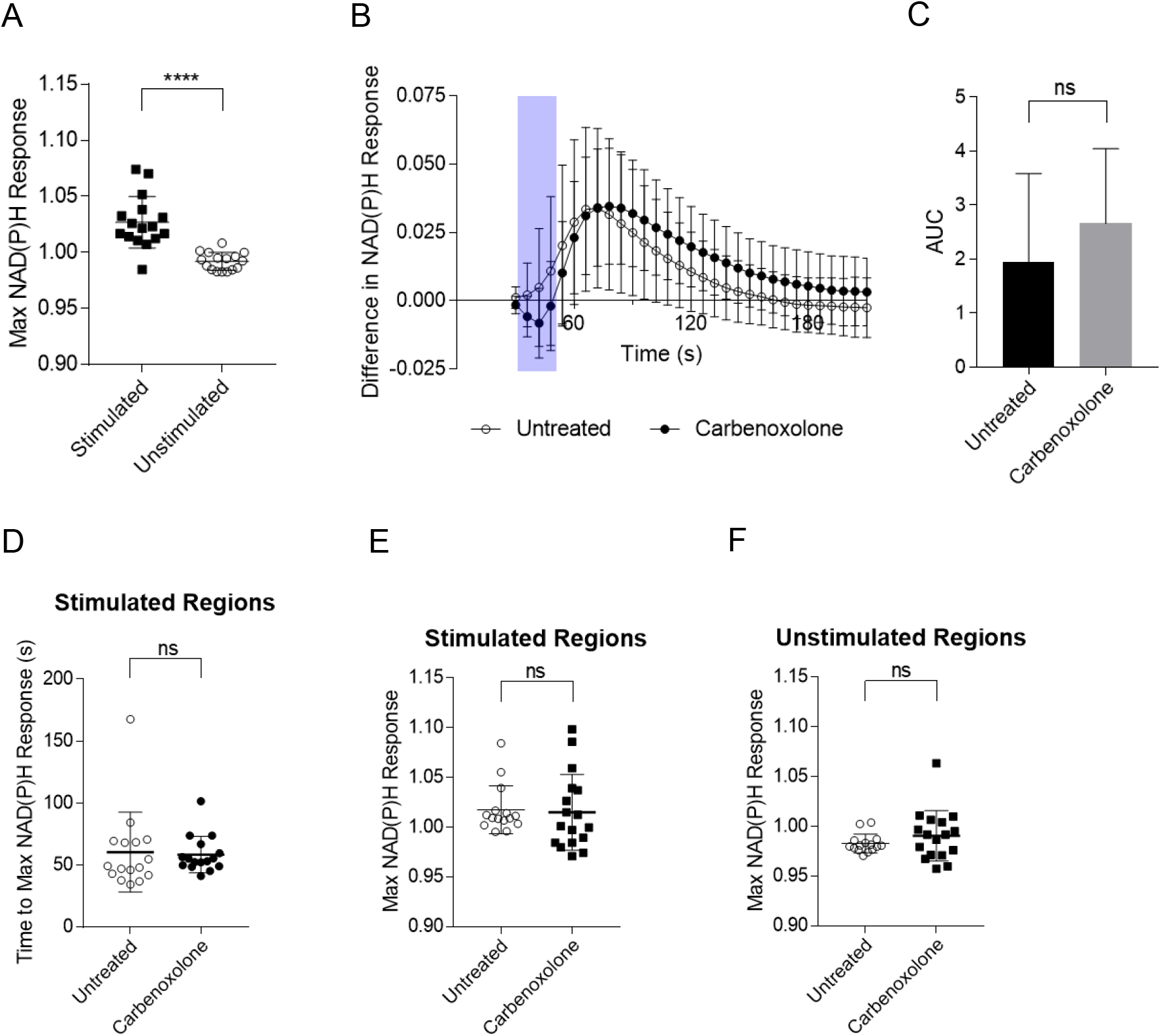
Coordination of metabolic response is consistent with diffusion of a molecule across gap junctions. (A) Islets were treated with 100 μM carbenoxolone, a gap junction inhibitor, before 20 mM glucose administration using a microfluidic applicator (n=16 islets). Max NAD(P)H responses are significantly greater in stimulated regions compared to unstimulated regions (p<0.0001). Error bars indicate S.D. (B) Treated islets were then compared to untreated islets. Glucose administration is indicated by the purple bar. The mean difference between responses in stimulated and unstimulated regions (symbols) ± S.D. (error bars) is depicted over time for untreated and carbenoxolone treatment (n=16 islets). (C) The mean area under the curve (AUC) ± S.D. for the curves in (B) is similar for both treatment conditions (n=16 islets, p=0.1985). (D) The time to max NAD(P)H responses in stimulated regions following glucose application is not altered by carbenoxolone treatment (n=16 islets, p=0.3608). (E, F) To test for off target effects, INS1E clusters, which lack gap junctions, were treated with carbenoxolone before glucose exposure (n=15 clusters). (E) Carbenoxolone treatment does not alter max NAD(P)H responses in stimulated (p=0.8215) or (F) unstimulated regions (p=0.2706) of INS-1E clusters. Means were compared by either student’s t-tests or matched pairs t-tests for max NAD(P)H response, AUC, and time to max NAD(P)H response.

Rises in intracellular calcium concentration may enhance metabolic responses^65,66^. Thus, it is possible that calcium regulates upstream metabolic processes (Figure 6A). To clarify the influence of cytoplasmic calcium concentration on metabolic responses, we activated ATP-sensitive potassium (KATP) channels, which trigger membrane depolarization, and blocked voltage gated calcium channels (VGCCs). Blockade of glucose-stimulated closure of KATP channels by diazoxide, or calcium influx by verapamil, significantly decreases islet calcium responses following glucose application (Supplemental Figure 2) (diazoxide: p=0.0104, p=0.0034; verapamil: p<0.0001, p<0.0001). NAD(P)H responses were not altered in stimulated or unstimulated regions (stimulated regions: p=0.1336, unstimulated regions: p=0.0522) by verapamil treatment (Figure 6B-C). However, diazoxide decreases NAD(P)H responses (p=0.0445) in islet cells outside the stimulation zone (Figure 6C). Therefore, NAD(P)H responses are independent of rises in cytoplasmic calcium, but may be influenced by membrane depolarization. Settings on the microfluidic applicator were held constant for all pharmacologic treatments, but the area and percent of the islet stimulated differed slightly between conditions (Supplemental Figure 3). This did not affect our interpretation of the data as neither the area nor the percent of the islet stimulated strongly correlated with NAD(P)H responses (Supplemental Figure 4).

**Figure 6.**
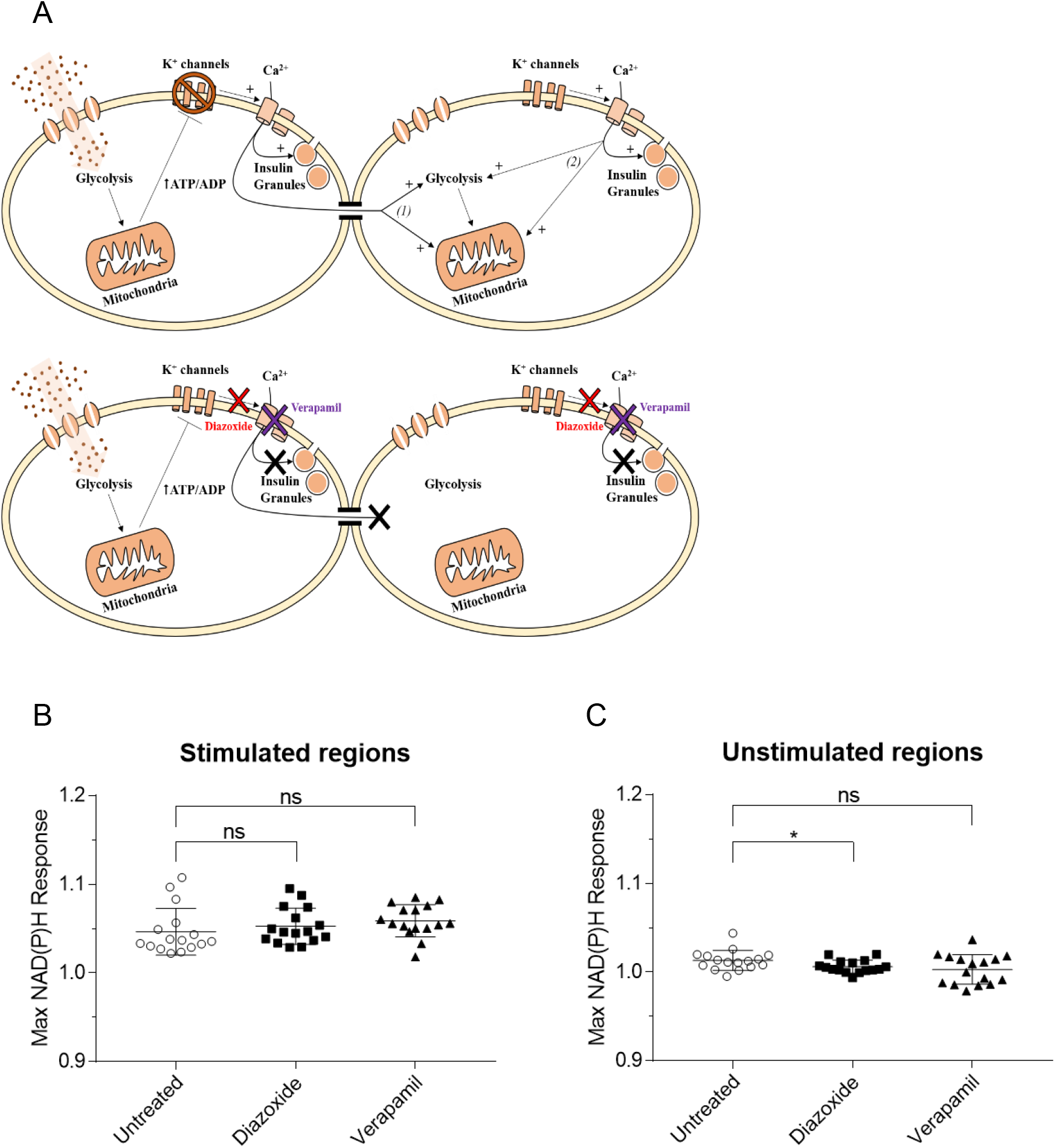
Metabolic responses of islets treated with an ATP-sensitive potassium channel (KATP) opener and a voltage gated calcium channel (VGCC) blocker following glucose stimulation. Islets were incubated with 250 μM diazoxide, a K_ATP_ opener, or 20 μM verapamil, a VGCC blocker before 20 mM glucose administration to portions of the islet. (A) The schematic diagram shows two connected beta cells. Glucose metabolism leads to closure of K_ATP_ channels and influx of calcium through voltage gated calcium channels in the left cell. Rises in metabolic activity in the right cell, which is not exposed to glucose, can occur through two potential calcium dependent mechanisms: (1) Calcium diffusion across gap junctions stimulates enzymes in glycolysis and the citric acid cycle. (2) Depolarization of the left cell triggers depolarization of the right cell. Then, calcium influx through VGCCs stimulates enzymes in glycolysis and the citric acid cycle. Preventing closure of K_ATP_ channels with diazoxide (red) or blocking VGCC with verapamil (purple) would inhibit calcium dependent rises in metabolic activity in cells not exposed to glucose. (B) Max NAD(P)H responses in stimulated regions are unchanged by treatment with diazoxide (n=16 islets, p=0.4645) or verapamil (n=16 islets, p=0.1336). (C) Max NAD(P)H responses in unstimulated regions are decreased by diazoxide (n=16 islets, p=0.0445), but are not affected by verapamil (n=16 islets, p=0.0522). Symbols denote individual islets. Mean ± S.D. is displayed. Significance was assessed by student’s t-tests.

Finally, we wanted to clarify the relationship between islet calcium and metabolic responses. As illustrated in Figure 3C-D, metabolic responses decay in magnitude and occur later in regions further from stimulation. Using the same regions of interest (ROIs), calcium responses were also measured. Rises in calcium occur approximately a minute after rises in NAD(P)H and are of comparable magnitude across the islet (Figure 7A). The similarity in the magnitude of calcium responses is also illustrated by comparing stimulated and unstimulated islet regions. Calcium responses are nearly identical in both regions (Figure 7B) in contrast to metabolic responses. Additionally, max calcium responses do not correlate with max NAD(P)H responses (Figure 7C-D). This is the case for integrated (area under the curve) calcium and NAD(P)H responses as well (Supplemental Figure 5). Taken together, our data suggest islet metabolic responses are coordinated via diffusion of metabolic intermediates across gap junctions, and this coordination is independent of rises in cytoplasmic calcium concentration.

**Figure 7.**
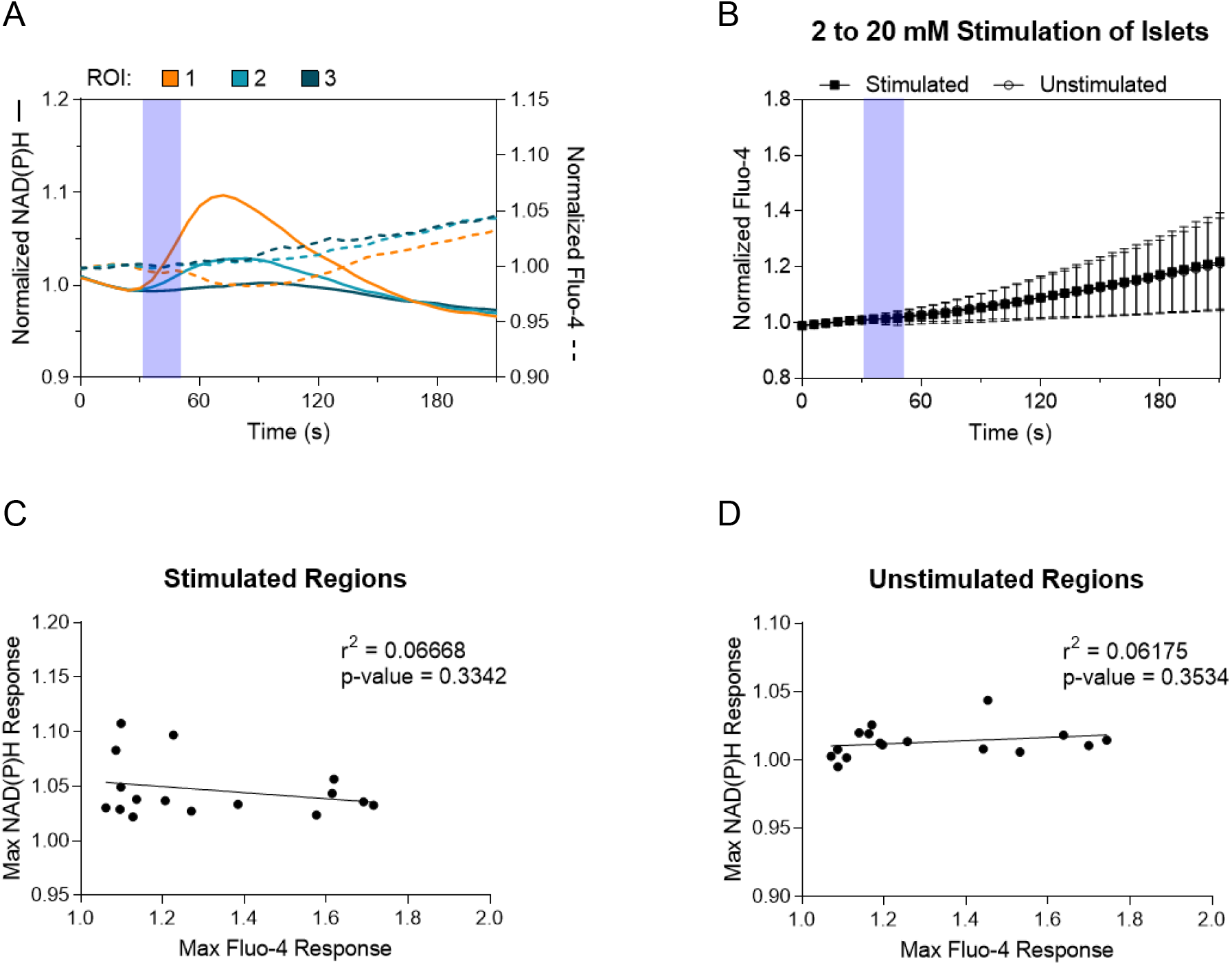
Metabolic responses are independent of rises in cytoplasmic calcium concentration. Calcium responses were measured using Fluo-4, a calcium indicator, following 20 mM glucose application, purple bars. (A) Calcium responses (dashed lines) of the regions of interest in Figure 3C are overlaid on top of metabolic responses (solid lines). (B) Mean calcium responses (symbols) ± S.D. (error bars) are shown for stimulated and unstimulated regions of islets (n=16 islets). Responses are almost identical between the two regions. Max NAD(P)H responses do not correlate with max Fluo-4 responses in stimulated (C) or unstimulated regions (D) of the islet (n=16 islets). Correlation was analyzed by linear regression.

## Discussion

Here we show that islet metabolic responses can spread from metabolically active cells to their neighboring beta cells. Following selective exposure of islet portions, cells located outside high glucose-stimulated areas displayed an increase in NAD(P)H. Gap junctional blockade inhibited this effect in islets, indicating that increased metabolism in unstimulated neighbors followed a transcellular pathway. Furthermore, metabolic activity did not spread among cells in INS-1E clusters. INS-1E cells lack gap junctions, and the metabolic responses in cells exposed to high glucose remained constrained by the spatial restriction of the BioPen. Given that these results reflected our observations in islets treated with carbenoxolone, we can reasonably conclude that gap junctions communicate metabolic responses across the islet.

Islet cell metabolic coupling also precedes calcium influx. Glucose-stimulated calcium responses follow metabolic responses by ~ 1 min. Unlike metabolic responses, calcium rises were synchronous throughout all islet cells, regardless of direct cellular stimulation with glucose. Additionally, glucose metabolism was unaffected by calcium channel inhibitors. Together, the data suggest that coordination of β-cell metabolism is calcium-independent. Based on the triggering pathway for insulin secretion and our evidence demonstrating that the signal occurs before VGCC activation, the mechanism likely involves a diffusing glycolytic product.

Diffusion of glycolytic intermediates most likely mediates metabolic communication between adjacent β-islet cells. Metabolic responses in untreated areas decayed linearly with distance and were delayed compared to the regions exposed to glucose (Fig 3). Further, NAD(P)H changes in areas outside the stimulation zone displayed decreased response amplitude and increasingly delayed peak response. In aggregate, the data show that the metabolic responses in unstimulated cells correlate with their distance from the stimulation edge. Maximum responses in unstimulated cells lagged over 10 s behind cells directly exposed to high glucose. Using the average length responses travel across the islet (30-50 μm), we roughly estimate a diffusion coefficient for the signaling molecules at around 60 – 180 μm^2^/s. These calculations place the molecular size of the communicating molecule at less than one kDa, assuming cytoplasmic diffusion^67–69^.

Glycolytic metabolites freely diffuse across the cytoplasm, unlike mitochondrial metabolism. Metabolite transport out of the mitochondria is comparatively slow and is thus an unlikely contributor to intra-islet communication. Size and charge constraints also need to be considered. The estimated pore size of connexin-36 gap junctions is between 15 and 45 angstroms^70,71^. As such, large molecules cannot diffuse across these gap junctions. Although connexin-36 gap junctions show a slight preference for cations^35,72^, anion permeation does occur^35,43,72–75^. Even so, the mechanism underlying connexin-36 gap junction selectivity and gating are not fully understood and may be regulated. Intracellular pH and magnesium ions can regulate conductance^76,77^ while glucose^78–80^, glibenclamide^34,73,78^ and IBMX^79^ can enhance gap junctional connectivity. Negatively charged metabolic intermediates should pass through islet gap junctions. Carboxyfluorescein (376 Da, charge = −2), an anionic dye similar in size and charge to glucose 6-phosphate (258 Da, charge = −2 freely diffuses across islet gap junctions^81^. Models of beta-cell signaling also demonstrate that glucose 6-phosphate diffuses through gap junctions^37^. From this, we propose that glucose 6-phosphate likely contributes to metabolic coordination in pancreatic islets.

A finer point illustrated by this study is that the molecule coordinating metabolic responses diffuses at least 30 μm. Previous studies of metabolite diffusion in neonatal rat islet monolayers show diffusion over similar lengths^36,38^. Our data show NAD(P)H increases in regions not exposed to glucose for up to 30 μm from the stimulation edge. Since the metabolite rise in the unstimulated areas is due to diffusion of a coordinating molecule, it follows that the molecule diffuses at least 30 μm. Metabolite diffusion may occur beyond this distance. However, data for all islets is unavailable.

Our findings support a role for pre-threshold metabolic coupling that is distinct from calcium-mediated glycolytic potentiation^66,82–84^. Previous models have proposed only post-action potential metabolic coupling. Coordination of metabolic responses is a consequence of synchronous rises in cytoplasmic calcium and occurs after insulin release. Our data suggest much earlier metabolic coupling. Calcium rises were identical in timing and magnitude in regions of the islet stimulated and unstimulated with glucose. Also, blocking VGCCs does not alter metabolic responses, as mentioned above. These results indicate that pre-threshold metabolic coupling is independent of glucose-stimulated calcium changes.

Interestingly, calcium responses do not peak within our experimental timeframe (within three minutes of stimulation). These data, along with the fact that calcium responses follow metabolic responses, open the possibility that metabolic coordination influences calcium coordination. The diffusion of glycolytic intermediates or other metabolites across the islet could increase the metabolic activity in a sufficient number of cells to trigger an electrical response producing increased calcium across the islet. Metabolic and calcium responses may not correlate in this scenario because membrane depolarization and activation of voltage-gated calcium channels may be an all or none rather than a graded response. Nevertheless, our data do not support or contradict this possibility.

KATP channel activity also affected the spread of metabolic activity across the islet. The summation of cellular KATP channel activity determines the electrical excitability of an islet^40^. Thus, membrane depolarization synchronizes across all β-islet cells, unlike metabolic activity, which we find linearly decays with distance. Yet, diazoxide decreases metabolic responses exclusively in unstimulated cells. This observation strengthens support for gap-junctionally mediated metabolic communication, given that hyperpolarization closes gap junctions^85–88^, including Cx-36^89^.

Metabolic-coupling may help explain the organization and potentiation of islet insulin secretion. While metabolic coupling agrees with earlier studies illustrating islet beta cells act as a syncytium^4,90^, it also allows for the possibility that beta cells with enhanced metabolic activity regulate islet metabolic and electrical activity. Notably, islet regions displaying high electrical control also have high metabolic activity^91^. Islet cell metabolic coupling, therefore, may be intrinsic to the mechanisms that control electrical coupling. In addition, the diffusion of molecules across coupled beta cells may enhance insulin secretion. Curiously, glucose can amplify insulin secretion independent of downstream effects on membrane potential or calcium concentration, which suggests glucose acts through alternate pathways to regulate insulin secretion^92^. Metabolite diffusion should add to the overall coupling mechanism since it occurs independently from action potentials and calcium oscillations. Thus, pre-threshold metabolic coupling likely contributes to the synergistic effect coupled beta cells have on glucose-stimulated insulin secretion^4–6^.

Johnston et al. recently proposed a new^41^, albeit controversial^93^ model for intra-islet β-cell communication. In their model, ‘hub’ or leader β-cells, enriched in glucokinase, drive calcium oscillations in follower beta cells. Our study supports metabolic communication between β-cells that flows from glycolytically more active to the less active cells. Given glucokinase’s role as a glucose sensor^12,94^, our studies imply that a cell with enhanced glucose phosphorylation would influence its neighbors. Even so, we find that gap junctions mediate pre-threshold metabolic communication. As a result, the need for direct cell-cell contact precludes the formation of networked activity as proposed^41,95^.

Type 2 diabetes mellitus progressively damages the ability of islets to secrete insulin over time. Given that loss or disruption of gap junction communication results in dysfunctional insulin secretion^28^, many have proposed a connection between gap-junctional connectivity and the decline of insulin secretion during diabetes^96^. For islet glucose metabolism, coupling defects may diminish islet metabolic activity and insulin secretion. Notably, decreased glucose-induced rises in NADH and ATP, as is observed with gap junction inhibition, impairs insulin secretion^55,97^. Further, damage to the most metabolically active cells, which are vulnerable to insults^98,99^ and may have disproportionate influence on islet activity, is a concern during type 2 diabetes. In turn, our findings suggest that human islets will be especially susceptible to the deterioration of metabolically active cells as beta cells are fewer in number compared to rodent islets^100^.

## Experimental procedures

### Islet isolation and culture

FVB mice were purchased from Jackson Labs, ME. Islets were isolated from 6-12 week old male and female FVB mice using collagenase XI (Sigma) digestion according to a previously published protocol^101^. Islets were cultured in DMEM growth media containing L-glutamine (Corning), 5.5 mM glucose, 10% fetal bovine serum (ThermoFisher), and 1% penicillin streptomycin solution (HyClone). Islets were seeded onto 50-mm dishes containing No. 1.5 glass coverslips (MatTek) coated with 0.01% poly-L-lysine (Sigma) for fluorescence imaging. Islets were imaged 5-7 days after seeding. All experiments were conducted in compliance with institutional guidelines and approved by the University of Maryland School of Medicine Institutional Animal Care and Use Committee.

### INS-1E cell culture

INS-1E cells were a gift from Dr. Peter Arvan^102^. Cells were cultured in RPMI-1640 growth media (Corning) containing L-glutamine, 11.1 mM glucose, 10% fetal bovine serum (ThermoFisher), 1% penicillin streptomycin solution (HyClone), 10 mM HEPES (Gibco), 1 mM sodium pyruvate (Gibco), and 50 μM β-mercaptoethanol (Sigma). Cells were passaged every 5 days using 0.25% trypsin-EDTA (Corning). To create clusters for fluorescence imaging, 50,000 cells were seeded onto 50-mm dishes containing No. 1.5 glass coverslips (MatTek) and maintained for 3-4 days prior to imaging.

### Microfluidic application of glucose

The Fluicell BioPen platform was used to selectively stimulate portions of clusters/islets with glucose. The BioPen tip was positioned ~50-75 μm away from the cluster/islet and 500 nM rhodamine 101 (Sigma) was used for visualization. Clusters/islets in all experiments were stimulated with glucose for 20 seconds using the following settings on the Fluicell BioPen platform: Pon = 280 mbar, Poff = 21, Vswitch = −115 mbar, Vrecirc = −115 mbar. All glucose solutions were made in imaging buffer (BMHH buffer containing 0.1% BSA).

### Fluorescence microscopy

Clusters/islets were incubated for 1 hr in imaging buffer (BMHH buffer containing 0.1% BSA and 2 or 5.5 mM glucose) at 37 °C prior to imaging. For calcium imaging, islets were similarly incubated for 1 hr in imaging buffer at 37 °C with the addition of 2.5 μM Fluo-4 (Invitrogen). Islets incubated with Fluo-4 were washed three times with imaging buffer prior to imaging. For experiments with pharmacologic treatments, cells/islets were treated with 100 μM carbenoxolone (Sigma), 20 μM verapamil (Sigma), or 250 μM diazoxide (Sigma) prior to stimulation. Images were collected using a Zeiss AxioObserver microscope equipped with a 20x, 0.8 NA Plan-Apochromatic objective and 1.25 Optovar tube lens. NAD(P)H autofluorescence was excited with a 365 nm LED and 49 DAPI filter cube. Fluo-4 fluorescence was excited with a 505 nm LED and 46 HE YFP filter cube. Images were collected at 37 °C using a Zeiss incubation system with a Zeiss Axiocam 506.

### Calculation of metabolic and calcium responses

Images were background subtracted and smoothed prior to calculation of responses. Responses were normalized to baseline and calculated in a pixel by pixel manner using a custom Python script. Max responses were determined from time points after stimulation was completed. Stimulated and unstimulated regions were determined based on thresholding of islets exposed to rhodamine. Islet and stimulation edges, responses based on distance, islet diameter, and time to max response were also determined using this script. For responses based on distance, the minimum distance between a given pixel and the islet/stimulation edge was used. The time to max response was calculated from only pixels exhibiting a response to glucose. Model Python code is available on GitHub (DOI: 10.5281/zenodo.4270289).

### Statistical analyses

All statistical analyses were performed using GraphPad Prism 7. Student’s t-tests were used to compare the mean responses. Matched pairs t-tests were used to compare responses from the same islet as was the case for responses in stimulated and unstimulated regions. Since imaging was not continuous, a Mann-Whitney or Wilcoxon matched-pairs signed rank test was used to compare time to max response. Finally, linear regression was used to evaluate the relationship between distance from islet/stimulation edge and NAD(P)H response, NAD(P)H response and area/percent stimulated, and max NAD(P)H response and max Fluo-4 response. Error bars in all figures denote standard deviation. 15 islet clusters (from three dishes) or 16 islets (from three mice, at least five islets per mouse) were imaged and analyzed for each experimental condition.

## Data availability

All data discussed are presented within the manuscript. Data will be shared upon request by contacting Vishnu Rao. The Python code used in this study is archived on Github (DOI: 10.5281/zenodo.4270289).

## Author contributions

V.P.R. collected all the data for the study, created all the figures, and drafted this manuscript. V.P.R. along with M.A.R. conceptualized the project and experimental design. M.A.R. served as the principal investigator for this study. She helped in drafting and editing of the manuscript as well as troubleshooting experiments.

## Funding and additional information

This work was funded by National Institutes of Health (NIH) Grants R01DK077140, R01MH111527, and R01MH111527 to M.R. NIH Training Grant T32GM008181 and NIH Fellowship Grant F30DK124986 supported V. P. R. The content is solely the responsibility of the authors and does not necessarily represent the official views of the National Institutes of Health.

